# Discovering dispersion: How robust is automated model discovery for human myocardial tissue?

**DOI:** 10.1101/2025.05.15.651144

**Authors:** Denisa Martonová, Sigrid Leyendecker, Gerhard A. Holzapfel, Ellen Kuhl

## Abstract

Computational modeling has become an integral tool for understanding the interaction between structural organization and functional behavior in a wide range of biological tissues, including the human myocardium. Traditional constitutive models, and recent models generated by automated model discovery, are often based on the simplifying assumption of perfectly aligned fiber families. However, experimental evidence suggests that many fibrous tissues exhibit local dispersion, which can significantly influence their mechanical behavior. Here, we integrate the generalized structure tensor (GST) approach into automated material model discovery to represent fibers that are distributed with rotational symmetry around three mean orthogonal directions—fiber, sheet, and normal—by using probabilistic descriptions of the orientation. Using biaxial extension and triaxial shear data from human myocardium, we systematically vary the degree of directional dispersion and stress measurement noise to explore the robustness of the discovered models. Our findings reveal that small dispersion in the fiber direction and arbitrary dispersion in the sheet and normal directions improve the goodness of fit and enable recovery of a previously proposed four-term model in terms of the isotropic second invariant, two dispersed anisotropic invariants and one coupling invariant. Our approach demonstrates strong robustness and consistently identifies similar model terms, even in the presence of up to 7% random noise in the stress data. In summary, our study suggests that automated model discovery based on the powerful generalized structure tensors is robust to noise and captures microstructural uncertainty and heterogeneity in a physiologically meaningful way.

## 1 Introduction

Computational modeling provides important insights into the intricate relationship between myocardial structure and mechanical function (Göktepe and Kuhl 2010; Baillargeon et al. 2014; Peirlinck et al. 2021; Martonová et al. 2022).

In particular, various passive material models have been proposed and investigated to accurately characterize the mechanical properties of myocardial tissue. These models aim to capture the complex anisotropic organization of the myocardium, which is critical for effective cardiac function.

In the myocardium, muscle fibers are arranged in a helical pattern, creating a three-dimensional architecture that supports the contraction and relaxation cycles of the heart. This arrangement allows for efficient twisting and squeezing motions necessary for effective blood pumping (Streeter et al. 1969; Holzapfel and Ogden 2009; Katz 2010; Holz et al. 2023). Myocardial fibers are arranged in layers of myocytes grouped together, so called sheets which exhibit a dynamic sliding behavior and contribute to ventricular deformation. Finally, to fully characterize the local orthotropic structure, we define the normal direction perpendicular to both fiber and sheet directions. Understanding this complex organization and possible uncertainties and anomalies in the microstructural architecture provide important insights into both normal physiology and pathological conditions. Traditional constitutive models, and recently discovered models based on the constitutive neural networks (Linka et al. 2023; Martonová et al. 2024; Peirlinck et al. 2024), often assume a perfect local alignment of a family of fibers at a given location, simplifying the fully anisotropic nature of the biological tissue, in particular the myocardium. This structural complexity necessitates the use of more advanced modeling techniques. Recent studies have explored how planar variations in fiber angles influence transversely isotropic model discovery for arteries (Vervenne et al. 2025), and studied similar effects in orthotropic textile structures (McCulloch and Kuhl 2024). Although these studies directly vary a single angle between two fiber families, various model frameworks exist to account for three-dimensional fiber dispersion, i.e., fibers do not have a fixed orientation at every location. One model framework was formulated by Lanir (1983), where each individual fiber within a dispersion is assigned a strain energy and the fibers are dispersed around a mean preferred direction according to an angular density distribution. This model framework was later modified (Sacks 2003; Driessen et al. 2005; Holzapfel and Ogden 2015; Martonová et al. 2021) and is referred to as angular integration (AI). The approach is based on full integration over a unit sphere, so that all possible fiber directions according to a given probability density function are considered. However, computational efficiency concerns have led to the adoption of alternative approximations such as the generalized structure tensor (GST) approach (Gasser et al. 2006; Holzapfel et al. 2015; Niestrawska et al. 2016) in which a GST is used. The advantages of this model framework include (i) that it is an algebraic formulation and therefore easier to implement than the AI formulation, (ii) it allows for explicit analytical results for a range of different deformations, (iii) the numerical analysis is less demanding, and (iv) it is more accurate because the numerical integrations required for the AI approach always lead to computational errors, whereas such integrations are not required for the GST model. The study of Holzapfel and Ogden (2017) documents that the predictive powers of the two models (GST, AI) are almost identical for a significant range of large deformations. A review on fiber dispersion modeling of soft biological tissues can be found by Holzapfel et al. (2019). In the present study, we utilize the GST approach and incorporate fiber, sheet, and normal dispersions into our constitutive neural network-based model discovery. In particular, instead of assuming a perfect fiber, sheet and normal alignment along one particular direction, we assume that these characteristic directions are locally distributed with some probability. Although the effects of myofiber dispersion on myocardial mechanics have been explored (Eriksson et al. 2013; Melnik et al. 2018; Guan et al. 2022), its impact on the discovery of material models remains insufficiently understood. Here we use dispersed invariants to explore the influence of uncertainty in the fiber architecture on model discovery. Additionally, to evaluate the impact of aleatoric noise, we introduce variability to the measured stress data. A key question we address is whether fiber dispersion and aleatoric noise result in the discovery of fundamentally different models or solely in modified parameter values.

We illustrate these effects through a case study on heart model discovery and compare the results with previously discovered models and parameters (Martonová et al. 2024) for given constant fiber, sheet, and normal orientations. To achieve this objective, we modify the architecture of the constitutive neural network to account for dispersions in the fiber, sheet, and normal directions and investigate the sensitivity of model discovery to both, the amount of the dispersion and the amount of noise in the stress-strain measurements. We train our network simultaneously on the biaxial extension and triaxial shear data (Sommer et al. 2015).

## 2 Methods

### 2.1 Continuum model

In continuum mechanics, a deformation map ***φ*** describes how material points move from their original (reference) configuration ***X*** to their current (deformed) configuration ***x*** = ***φ***(***X***) (Gurtin 1981; Holzapfel 2000). The deformation gradient **F** of the map ***φ*** with respect to the undeformed coordinates ***X*** and its determinant *J* are defined as

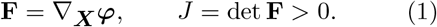

Multiplying the deformation gradient with its transpose **F**^t^ from the left, we obtain the right Cauchy-Green deformation tensors **C** = **F**^t^ ·**F**. We demonstrate our approach using human myocardial tissue, modeled as perfectly incompressible orthotropic material with three distinct structural directions. As illustrated in Figure 1 right, these directions correspond to the fiber, sheet, and normal orientations, denoted by ***f*** _0_, ***s***_0_ and ***n***_0_, respectively. We introduce nine invariants to describe the deformation (Spencer 1984; Holzapfel and Ogden 2009), three standard isotropic invariants *I*_1_, *I*_2_, *I*_3_, three anisotropic invariants characterizing the stretches, *I*_4f_, *I*_4s_, *I*_4n_, and three mixed coupling invariants, *I*_8fs_, *I*_8fn_, *I*_8sn_,

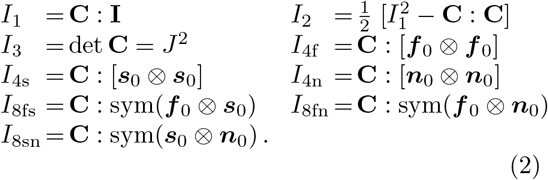

Under our assumption of perfect incompressibility, the third invariant equals one, *I*_3_ = *J* ^2^ = 1.

**Fig. 1.**
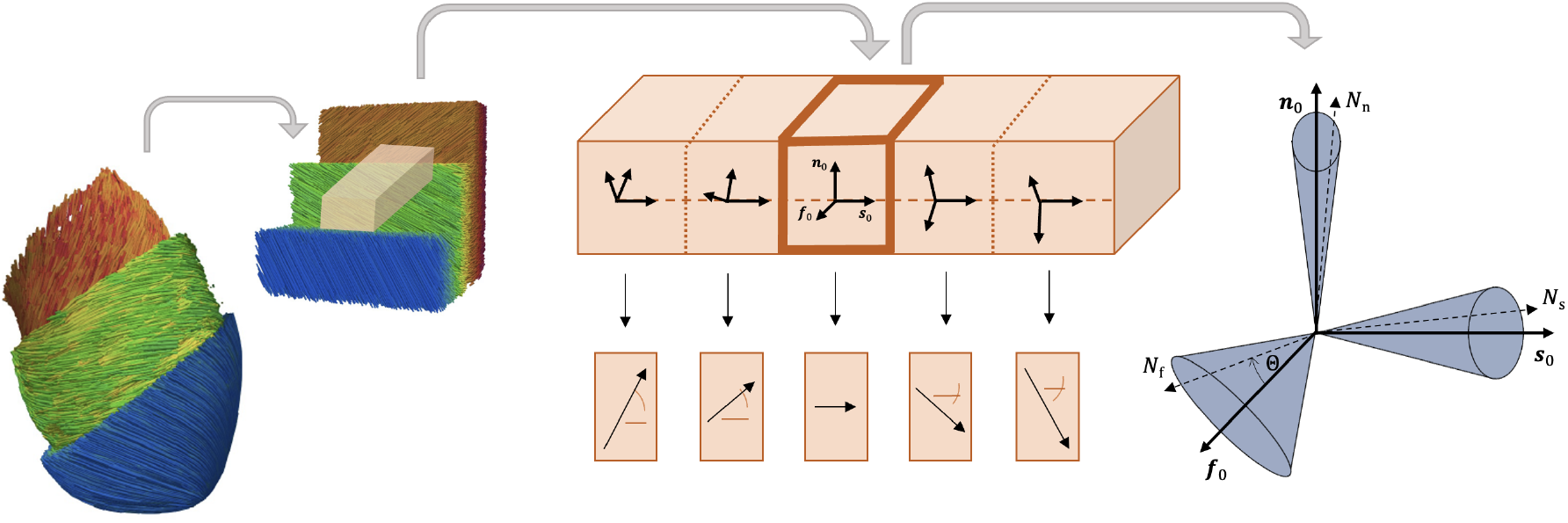
Fiber, sheet and normal architecture in the left ventricle. Left and middle: Schematic representation of fiber architecture in left ventricle, rotating throughout the ventricular wall. Right: Schematic representation of the dispersion local in fiber, sheet, and normal directions. Fibers, sheets and normals are assumed to be located inside the blue cone.

### 2.2 Material model discovery with constitutive neural networks

To autonomously discover the most suitable passive material model for human myocardial tissue with probabilistic fiber, sheet and normal orientations, we use constitutive neural networks as our model discovery framework (Linka et al. 2021; Linka and Kuhl 2023). In particular, we select an orthotropic constitutive neural network made up of two hidden layers and 32 nodes with built-in incompressibility and polyconvexity (Martonová et al. 2024). As shown in Figure 2, eight invariants serve as network inputs and the network output is a single scalar-valued free-energy function *ψ*. This network architecture is therefore capable of discovering up to 2^32^ possible models. In the present work, we include uncertainty in both network input and network output, to better reflect realistic experimental data.

**Fig. 2.**
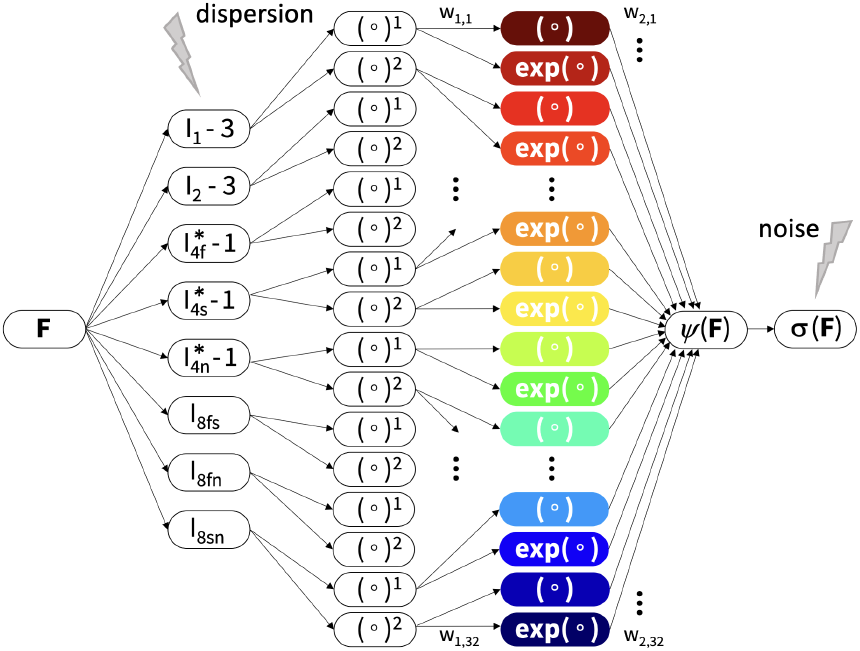
Orthotropic, perfectly incompressible constitutive neural network accounting for fiber, sheet and normal dispersion. Eight invariants 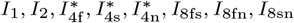 serve as network input, while a scalar-valued free energy function *ψ*, depending on these eight invariants, is the network output. Cauchy stress components are derived from the discovered free energy function *ψ*. The displayed feed-forward partially connected network contains two hidden layers. In the first layer, the corrected eight invariants are raised to the first and second powers, (∘) and (∘)^2^, while the identity (∘) and exponential functions (exp(∘) operate on these values.

#### 2.2.1 Network input uncertainty – fiber dispersion

We follow the GST approach (Gasser et al. 2006) to include fiber, sheet and normal dispersion into our network. This approach is based on a symmetric GST **H**_*i*_ for each fiber family *i*, defined as

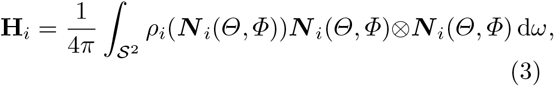

where 𝒮 ^2^ = {***N*** _*i*_ : |***N*** _*i*_| = 1} is the unit sphere, d*ω* = sin d*Θ* d*Φ* and *ρ*_*i*_(***N*** _*i*_(*Θ, Φ*)) is an orthotropic probability density function and ***N*** _*i*_ is any unit vector in three-dimensional Eulerian space, i.e. ***N*** _*i*_ denotes a possible fiber direction within fiber family *i*, and *ρ*_*i*_(***N*** _*i*_(*Θ, Φ*))d*ω* represents the normalized number of fibers with orientations within the intervals [*Θ, Θ* +d*Θ*], [*Φ, Φ*+d*Φ*] that fulfills the symmetry condition *ρ*_*i*_(***N*** _*i*_) = *ρ*(**−*N*** _*i*_) (Gasser et al. 2006).

Now, we assume axisymmetric distributions for each family of undeformed fibers with the mean direction ***i***_0_. The density function is then independent of *Φ* and simplifies to *ρ*_*i*_(***N*** _*i*_(*Θ, Φ*)) ≈*ρ*_*i*_(*Θ*). For a given family of fibers, the fibers are distributed with rotational symmetry about a mean referential direction represented by a unit vector ***i***_0_, the GST approach in Eq. (3) simplifies to

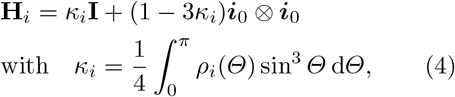

where **I** is the identity tensor and *κ*_*i*_ ∈ [0, 1*/*3] is the dispersion parameter. In the following, we consider three fiber families, fibers, sheets, and normals, *i* ∈ {f, s, n}, and we assume that the orientations of these fiber families follow a modified *π*-periodic von Mises distribution. The probability density function centered at *Θ* = 0 is then given by

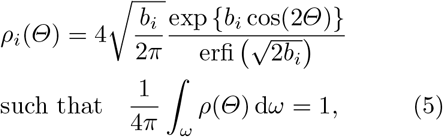

with a concentration parameter *b*_*i*_ *>* 0 and an imaginary error function erfi(*x*) = −*i* erf(*x*), with

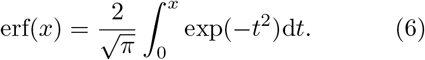

We can interpret this distribution as a projection of the normal distribution onto the unit sphere (Fisher et al. 1987; Gasser et al. 2006). In particular, for *κ*_*i*_ = 0, we recover a perfect fiber alignment along the mean direction ***i***_0_ whereas for *κ*_*i*_ = 1*/*3, the fibers are distributed isotropically within the sphere, as schematically shown in Figure 3. Using the GST approach, we modify the three anisotropic invariants *I*_4f_, *I*_4s_, *I*_4n_, and obtain the following dispersed invariants for our network input,

**Fig. 3.**
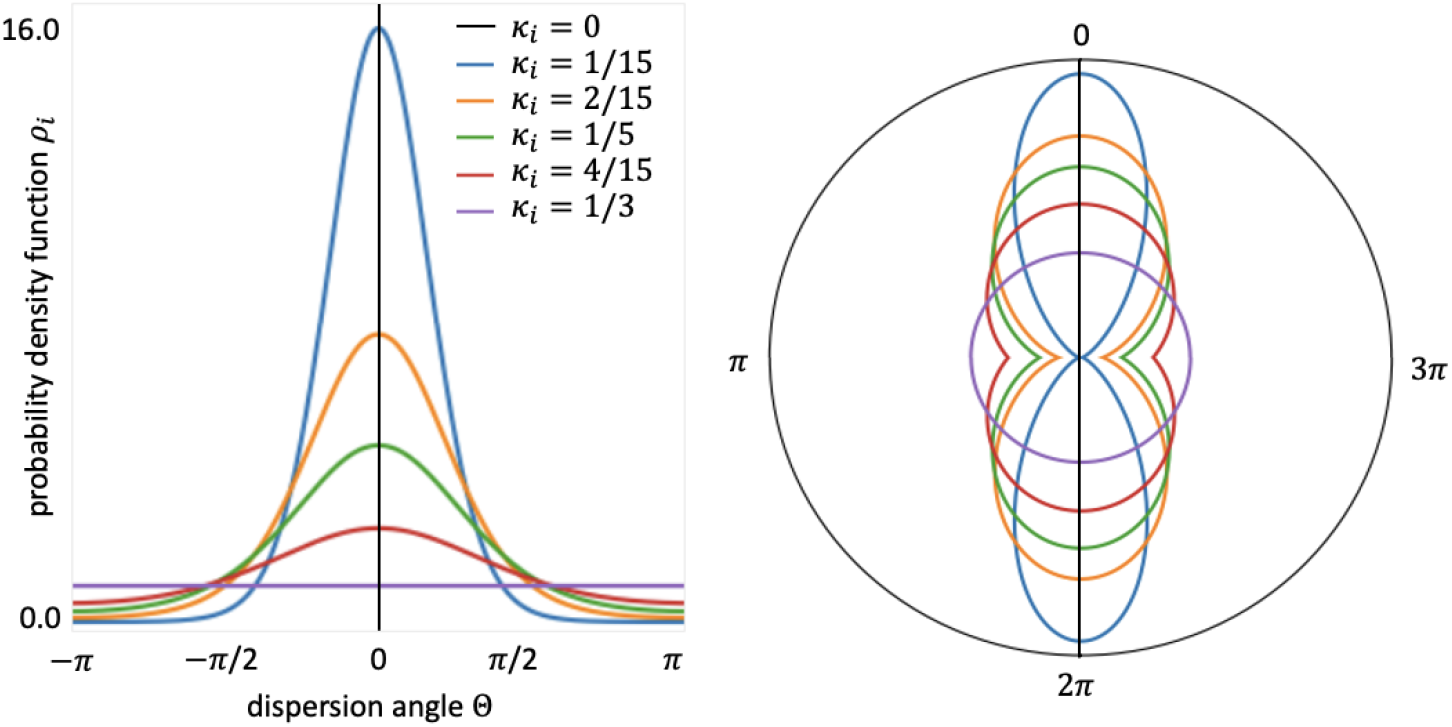
Modified von Mises probability density function for the dispersion angle *Θ*. Left: Two-dimensional probability density function *ρ*_*i*_ for different dispersion parameters *κ*_*i*_. Right: Planar schematic representation of possible fiber, sheet and normal directions for a given dispersion parameter *κ*_*i*_.

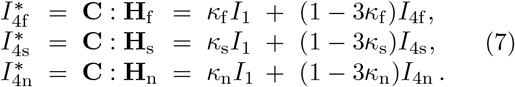

Following previous studies (Eriksson et al. 2013; Guan et al. 2022), we solely account for the dispersion in the fourth invariants. We note, though, that it is also possible to account for dispersion in the modified coupling *I*_8_-like invariant, see (Melnik et al. 2018). However, the softening effect of this coupled dispersed invariant was shown to be smaller than for the dispersed fourth invariants and its physical interpretation remains largely unexplored.

#### 2.2.2 Network output uncertainty – added Gaussian noise

To investigate the influence of aleatoric uncertainty on the discovered model and its parameters, we systematically add varying levels of Gaussian noise, *n*_*k*_, *k* ∈ {0.03, 0.05, 0.07, 0.1}, to the measured values of the Cauchy stress components *σ*_ij_, i.e.

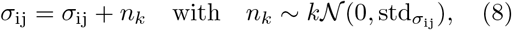

where 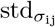 is the standard deviation of the measured Cauchy stresses for a given deformation mode.

#### 2.2.3 Neural network architecture

Figure 2 showcases the architecture of our neural network with its two hidden layers and 32 nodes. The network assumes perfect incompressibility and takes eight input invariants: two isotropic invariants *I*_1_ and *I*_2_; three anisotropic invariants that account for fiber, sheet, and normal dispersion 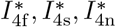; and three coupling invariants *I*_8fs_, *I*_8fn_, *I*_8sn_. The output is a scalar-valued free energy function *ψ*. In the first layer, the network generates the powers (∘) and (∘)^2^ of the corrected input invariants. In the second layer, the network applies the identity function (∘) and the exponential function (exp(∘)) to these values. This allows us to explicitly express the free energy function *ψ*,

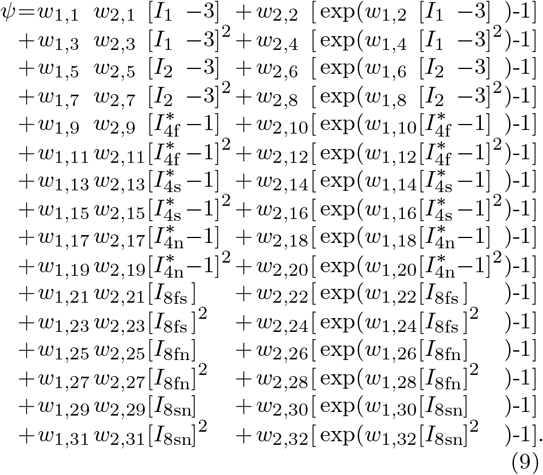

Due to the incompressibility assumption, the free energy is modified by the hydrostatic pressure term *p*, yielding *ψ* = *ψ* − *p* [*J* − 1]. The corrections for the invariants values by one and three ensure that *ψ*(**F** = **I**) = 0 is satisfied. We note that we only activate the anisotropic dispersed fourth invariants, 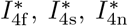, if their directions are under tension (Holzapfel and Ogden 2009). For the eighth coupling invariants, *I*_8fs_, *I*_8fn_, *I*_8sn_, in the undeformed configuration, the values are zero and can be used as such. Notably, these coupling invariants are sign-sensitive with respect to the fiber, sheet, and normal directions and are therefore not strictly invariant (Holzapfel and Ogden 2009; Melnik et al. 2018). However, when training the network with experiments specified in Section 2.3, the sign always remains positive. Following standard arguments of thermodynamics, we obtain the Cauchy stress from the free-energy function *ψ* in Eq. (9) as

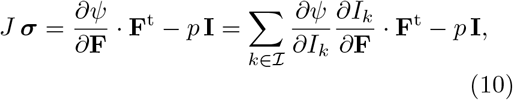

where 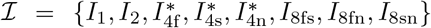 is the set of network input invariants. Analytical expressions for the free-energy derivatives with respect to the eight invariants depend on the network weights and are documented in prior work (Martonová et al. 2024). The derivatives of these invariants with respect to the deformation gradient follow as

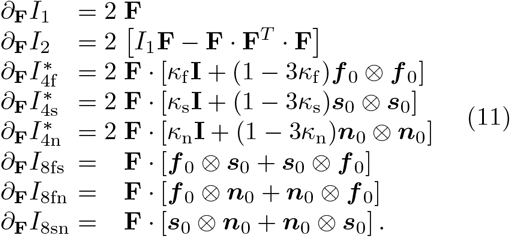

### 2.3 Mechanical experiments used for training

Finally, we evaluate the stresses for the specific experimental loading modes. Motivated by our previous findings (Martonová et al. 2024), we train our dispersed-invariant network from Figure 2 simultaneously with triaxial shear and biaxial extension data from human myocardial tissue (Sommer et al. 2015). In the following, the subscripts, f, s, n are associated with the fiber, sheet, and normal directions, respectively, and are used to denote the corresponding stretches *λ*_i_ and shear strains *γ*_ij_, i, j ∈ {f, s, n}. For all six triaxial shear experiments, the three stretches remain constant and equal to one, i.e. *λ*_f_ = *λ*_s_ = *λ*_n_ ≡ 1. During triaxial shear testing in the ij-plane along the i-direction, only the shear strain *γ*_ij_ becomes nonzero, while all other shear strains remain zero. As a result, each of the six tests yields two nonzero shear stress components, *σ*_ij_ = *σ*_ji_ ≠ 0, in particular

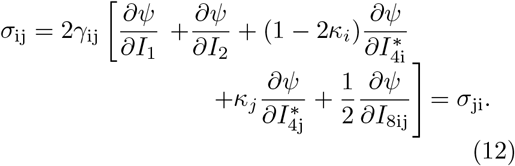

We note that due to the fiber, sheet, and normal dispersions, *κ*_*i*_ *>* 0, both fourth invariants, 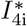 and 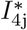, contribute to the shear stress *γ*_ij_, whereas in the non-dispersed case, *κ*_*i*_ = *κ*_*j*_ = 0, only the invariant 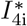 appears in Eq. (12).

For the biaxial extension tests, we consider five different ratios of fiber and normal stretches (*λ*_f_ ≥ 1) : (*λ*_n_ ≥ 1), namely 1:1, 1:0.5, 1:0.75, 0.5:1, 0.75:1. The remaining sheet stretch is computed from the incompressibility condition as *λ*_s_ = 1*/*[*λ*_f_ *λ*_n_] ≤ 1 and all shear strains vanish. We further assume a zero stress condition throughout the sample thickness, such that the condition *σ*_ss_ = 0 holds. For the hydrostatic pressure *p* in (10), we obtain

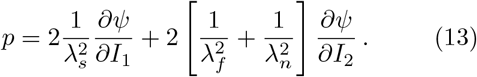

The non-zero normal stresses take the following form,

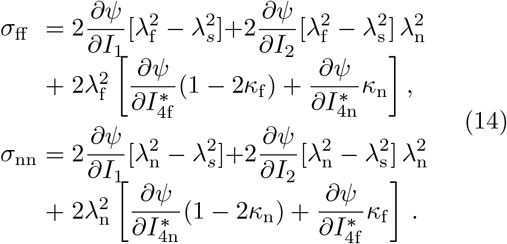

In Eqs. (12) and (14), the free energy derivatives with respect to the eight invariants depend on the network weights (Martonová et al. 2024), which we learn during neural network training.

### 2.4 Neural network training

Using the explicit formulations for all Cauchy stress components, we train the network by optimizing a loss function *L* (see Eq. (15) below), which penalizes the mean squared error in the *L*_2_-norm between the modeled Cauchy stress ***σ***(**F**_*i*_, **w**) and the measured Cauchy stress 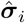, divided by the total number of data points *n*_data_. Eq. (9) suggests that we can replace *w*_i,1_ and *w*_i,2_ by their product *w*_i,1_*w*_i,2_ for all odd-indexed weights *i* = 2*n* + 1, *n* ≥ This reduces the total number of non-negative trainable weights from 64 to 48, **w** = {*w*_1,1_*w*_2,1_, *w*_1,2_, *w*_2,2_, …, *w*_1,32_, *w*_32,2_} ≥ **0**. The first-layer weights *w*_1,j_ are unit-less parameters and the second-layer weights *w*_2,j_ have units of the stiffness. To promote model sparsity and enhance model interpretability, we add a *L*_1_- regularization term *α* ∥**w**∥ _1_, to our loss function (McCulloch et al. 2023), leading to

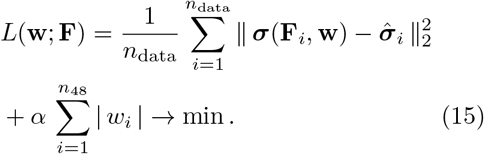

In the following, we set the regularization parameter *α* = 0.01, as proposed in recent studies (Martonová et al. 2024; Vervenne et al. 2025). To minimize the loss function in Eq. (15), we use the adaptive first-order gradient-based optimizer Adam (Kingma and Ba 2014). The network is trained for up to 30 000 epochs with a batch size of 32. We implement early stopping criterion if accuracy does not improve for 1 000 consecutive epochs. To mitigate the risk of convergence to local minima, we initialize the network weights randomly. Specifically, we use the Glorot normal initializer for layers with identity functions, and a random uniform initializer for layers with exponential functions, assigning non-negative weights with a maximum value of 0.1. We quantify model performance using the coefficient of determination R^2^, both individually for each individual experiment and combined across all experiments.

## 3 Results and Discussion

To systematically investigate the influence of the input and output noise, we perform the following learning scenarios:

- we vary the amount of the Gaussian noise added the experimental data according to Eq. (8) and we assume perfect fiber, sheet and normal alignment (*κ*_f_ = *κ*_s_ = *κ*_n_ = 0),
- we add 3% Gaussian noise to the experimental data and vary the dispersion in the fiber direction (*κ*_f_≠ 0); we assume perfect alignment in sheet and normal directions (*κ*_s_ = *κ*_n_ = 0),
- we add 3% Gaussian noise to the experimental data and we vary the dispersion in the sheet and normal directions (*κ*_s_ = *κ*_n_≠ 0); we assume perfect alignment in fiber direction (*κ*_f_ = 0),
- we add 3% Gaussian noise to the experimental data and we vary the dispersion in the fiber, sheet and normal directions (*κ*_f_ = *κ*_s_ = *κ*_n_≠ 0).

### The network robustly discovers four-term models, even with aleatoric noise on the experimental data

The robustness of our model discovery process is demonstrated by its insensitivity to the introduced random noise according to Eq. (8). As shown in Figure 4, up to the data perturbations of 7% random Gaussian noise, our constitutive neural network consistently identifies similar four-term models and eight key parameters, with only minor variations. At the chosen regularization level of *α* = 0.01, the invariants 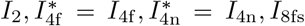 contribute quadratically to the free energy *ψ* and are highlighted in orange, yellow, turquoise, and blue in Figure 4. Notably, the data noise of 10% activates an additional invariant 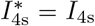. Interestingly, this term is also present in the classical Holzapfel Ogden model for cardiac tissue (Holzapfel and Ogden 2009).

**Fig. 4.**
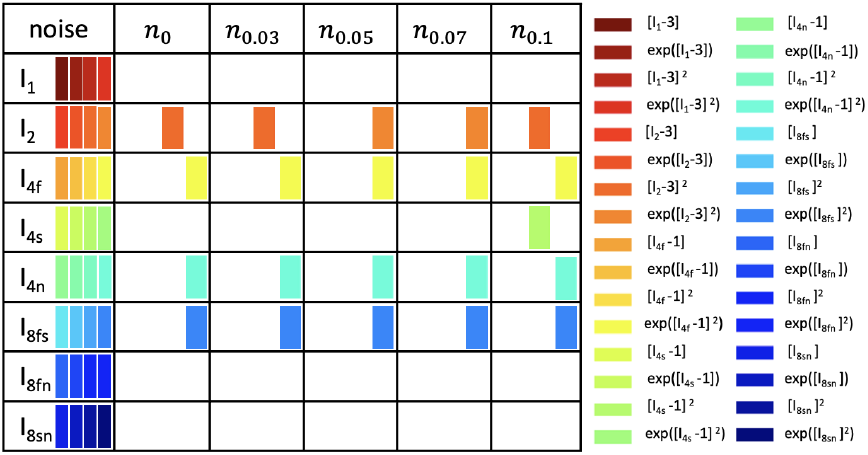
Model discovery for human myocardium with varying Gaussian noise. Active invariants for varying amount of Gaussian noise, 0%,3%,5%,7%,10%, added to the measured stress components according to Eq.(8); color-coded blocks represent the contributions of the particular active term out of four possible terms, depending on one of the eight possible invariants given in the first column.

### The network robustly discovers four-term models, even with dispersion in sheet and normal directions

As highlighted in Figure 5 left, for all variations of the dispersion parameters *κ*_s_ and *κ*_n_, we recovered four-term models which depend quadratically on the four invariants 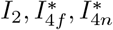 and *I*_8*fs*_. The plot in Figure 5 right indicates that, with increasing dispersion, the stiffness in the normal direction, quantified by the product of the network weights *w*_1,20_ *w*_2,20_, increases. This is reasonable, since a dispersed normal direction will result in reduced contributions to the strain energy in the fn-plane during biaxial extension and in the fn- and sn-planes during shear testing. Interestingly, as suggested by the last row in Table 1, the mean goodness of fit slightly improves, from R^2^ = 0.890 for *κ*_f_ = *κ*_s_ = *κ*_n_ = 0 to R^2^ = 0.922 for *κ*_s_ = *κ*_n_ = 4*/*15.

**Table 1.** Discovered material parameters for human myocardial tissue for varying amount of sheet and normal dispersions. Interpretable network weights for simultaneous training with six shear and five biaxial tests with a regularization parameter *α* = 0.01 for six different pairs of dispersion parameters *κ*_s_ = *κ*_n_ ∈ {0, 1*/*15, 2*/*15, 1*/*5, 4*/*15, 1*/*3}, *κ*_f_ = 0; mean goodness of fit R^2^.

| network weights | model term | $\kappa_s = \kappa_n$ | | | | | |
| --- | --- | --- | --- | --- | --- | --- | --- |
| $w_{1,\bullet}, [-], w_{2,\bullet}$ [kPa] | $[-]$ | 0 | 1/15 | 2/15 | 1/5 | 4/15 | 1/3 |
| $w_{1,7} \cdot w_{2,7}$ | $[I_2 - 3]^2$ | 5.153 | 5.108 | — | 5.144 | — | — |
| $w_{1,8}, w_{2,8}$ | $\exp([I_2 - 3]^2)$ | — | 1.075, 4.520 | — | — | 1.215, 3.813 | 1.401, 2.572 |
| $w_{1,12}, w_{2,12}$ | $\exp([I_{4f}^* - 1]^2)$ | 21.062, 0.081 | 23.844, 0.063 | 23.331, 0.065 | 23.925, 0.059 | 24.687, 0.055 | 24.214, 0.055 |
| $w_{1,20}, w_{2,20}$ | $\exp([I_{4n}^* - 1]^2)$ | 4.132, 0.340 | 4.683, 0.380 | 3.565, 0.705 | 2.598, 1.358 | 2.759, 2.228 | 4.044, 4.202 |
| $w_{1,23} \cdot w_{2,23}$ | $[I_{8fs}]^2$ | — | 0.257 | — | — | — | — |
| $w_{1,24}, w_{2,24}$ | $\exp([I_{8fs}]^2)$ | 0.511, 0.485 | — | 0.517, 0.532 | 0.537, 0.514 | 0.605, 0.607 | 1.696, 0.351 |
| $R^2$ | | 0.890 | 0.898 | 0.905 | 0.914 | 0.922 | 0.920 |

**Fig. 5.**
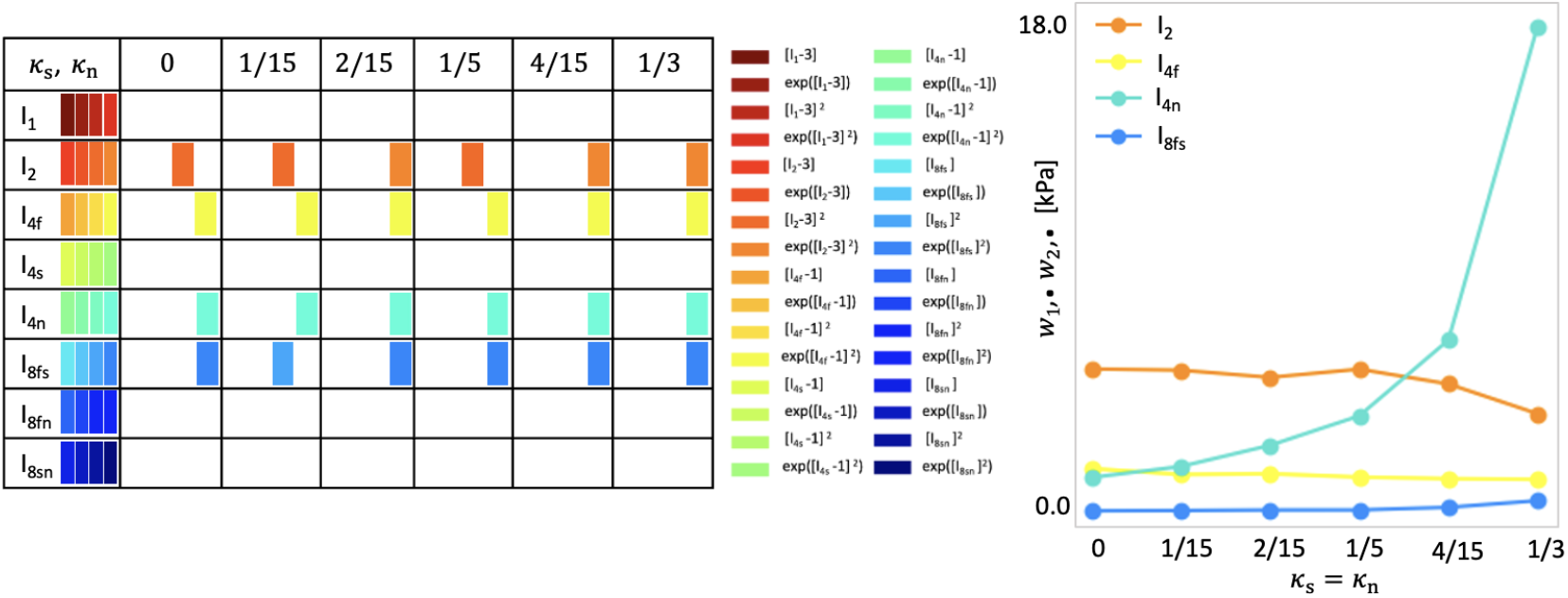
Model discovery for human myocardium with with varying amount of dispersion in sheet and normal directions. Left: Active invariants for varying amount of dispersion in sheet and normal directions, *κ*_s_ = *κ*_n_ ∈ {0, 1*/*15, 2*/*15, 1*/*5, 4*/*15, 1*/*3}, *κ*_f_ = 0; color-coded blocks represent the contributions of the particular active term out of four possible terms, depending on one of the eight possible invariants given in the first column. Right: Stiffness-like weight values for active invariants; the sum of products 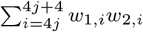, where *j* = 1, *j* = 2, *j* = 4, *j* = 5 for contributions of terms depending on *I*_2_, *I*_4f_, *I*_4n_, *I*_8fs_.

### Model discovery is robust against small dispersion in fiber direction

Figures 6 and 7 showcase that, for dispersion up to *κ*_f_ = 1*/*5 with perfectly aligned sheet and normal directions and for the dispersion up to *κ*_f_ = *κ*_s_ = *κ*_n_ = 2*/*15, we recover four-term models which depend on the four invariants 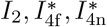 and *I*_8fs_. Moderate and high dispersions in fiber direction induce model sparsity and reduces the goodness of fit. With increasing dispersion in fiber direction, we observe an increasing stiffness of the fiber, measured by the product *w*_1,12_ *w*_2,12_, suggested by the yellow curve in Figures 5 and 7 right. This nearly exponential increase can be explained by the softening effect of fiber dispersion (Melnik et al. 2018). In the extreme case of fully dispersed fibers (*κ*_f_ = 1*/*3), the fiber contribution is spread out over the entire three-dimensional sphere as visualized in Figure 3 right. Figures 8 and 9 depict the individual contributions of specific model terms to the modeled stress components. It is obvious that the assumed dispersion in fiber and sheet directions activates the invariant 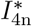 in the modeled fiber stress *σ*_ff_, see second row in the respective figures. Analogously, the invariant 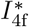 is activated when modeling the normal stress *σ*_nn_, see third row in the respective Figures. For the full dispersion *κ*_f_ = *κ*_s_ = *κ*_n_, both invariants, 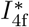 and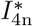, contribute to the overall stress equally resulting in the equal modeled stresses *σ*_nn_ = *σ*_nn_. We note that rescaling the stress-axis in the second and third rows in Figure 9 would result in the equal plots rows two and three. However, this equality does not correspond to the experimental data, which is as well reflected by the low R^2^ = 0.722 given Table 2.

**Table 2.**
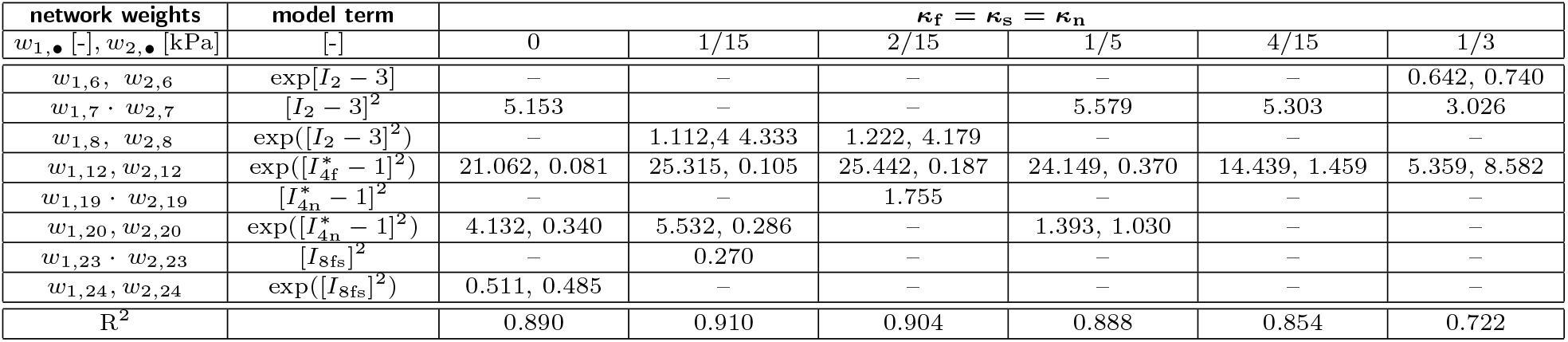
Discovered material parameters for human myocardial tissue for varying amount of fiber, sheet and normal dispersions. Interpretable network weights for simultaneous training with six shear and five biaxial tests with a regularization parameter *α* = 0.01 for six different triples of dispersion parameters *κ*_f_ = *κ*_s_ = *κ*_n_ ∈{0, 1*/*15, 2*/*15, 1*/*5, 4*/*15, 1*/*3}; mean goodness of fit R^2^.

### Across all dispersion settings, the network consistently discovers models depending on the second invariant

Interestingly, all discovered models for all levels of dispersion incorporate the second invariant rather than the first. As shown in Tables 1, 2 and 3, except for the full dispersion in the fiber direction, the discovered models rely exclusively on the quadratic term in the second invariant *I*_2_. Notably, as visualized in Figures 5, 6 and 7, the product of the discovered weights *w*_1,7_ *w*_2,7_ for the solely quadratic term and *w*_1,8_*w*_2,8_ for the quadratic exponential term changes only minimally, with its minimum of 4.63 kPa for *κ*_s_ = *κ*_n_ = 4*/*15 and maximum of 5.58 kPa *κ*_f_ = *κ*_s_ = *κ*_n_ = 1*/*5. This selective preference is visually corroborated in Figures 8 and 9, where the color-coded stress terms are dominated by orange contributions from the second invariant *I*_2_. The striking influence of the second invariant contrasts with the commonly used models that depend solely on the first invariant (Treloar 1948; Demiray 1976; Lanir 1983; Holzapfel and Ogden 2009; Budday et al. 2017; Guan et al. 2019), but it aligns well with a few previous models (Weiss et al. 1996; Horgan and Smayda 2012) and with recently discovered models (Kuhl and Goriely 2024; Linka and Kuhl 2024; Martonová et al. 2024; Vervenne et al. 2025) for soft biological tissues.

**Table 3.** Discovered material parameters for human myocardial tissue for varying amount of fiber dispersion. Interpretable network weights for simultaneous training with six shear and five biaxial tests with a regularization parameter *α* = 0.01 for six different dispersion parameters *κ*_f_ ∈ {0, 1*/*15, 2*/*15, 1*/*5, 4*/*15, 1*/*3}, *κ*_s_ = *κ*_n_ = 0; mean goodness of fit R^2^.

| network weights | model term | $\kappa_f$ | | | | | |
| --- | --- | --- | --- | --- | --- | --- | --- |
| $w_{1,\bullet}, [-], w_{2,\bullet}$ [kPa] | $[-]$ | 0 | 1/15 | 2/15 | 1/5 | 4/15 | 1/3 |
| $w_{1,6}, w_{2,6}$ | $\exp[I_2 - 3]$ | — | — | — | — | — | 0.629, 0.721 |
| $w_{1,7} \cdot w_{2,7}$ | $[I_2 - 3]^2$ | 5.153 | 5.121 | 5.290 | 5.527 | 5.308 | 3.080 |
| $w_{1,12}, w_{2,12}$ | $\exp([I_{4f}^* - 1]^2)$ | 21.062, 0.081 | 25.558, 0.105 | 24.893, 0.199 | 23.176, 0.406 | 14.619, 1.438 | 5.059, 9.141 |
| $w_{1,19} \cdot w_{2,19}$ | $[I_{4n}^* - 1]^2$ | — | — | — | 0.608 | — | — |
| $w_{1,20}, w_{2,20}$ | $\exp([I_{4n}^* - 1]^2)$ | 4.132, 0.340 | 5.517, 0.214 | 5.644, 0.161 | — | — | — |
| $w_{1,21} \cdot w_{2,21}$ | $[I_{8fs}]$ | — | — | 0.104 | 0.035 | — | — |
| $w_{1,23} \cdot w_{2,23}$ | $[I_{8fs}]^2$ | — | 0.222 | — | — | — | — |
| $w_{1,24}, w_{2,24}$ | $\exp([I_{8fs}]^2)$ | 0.511, 0.485 | — | — | — | — | — |
| $R^2$ | | 0.890 | 0.905 | 0.904 | 0.885 | 0.852 | 0.720 |

**Fig. 6.**
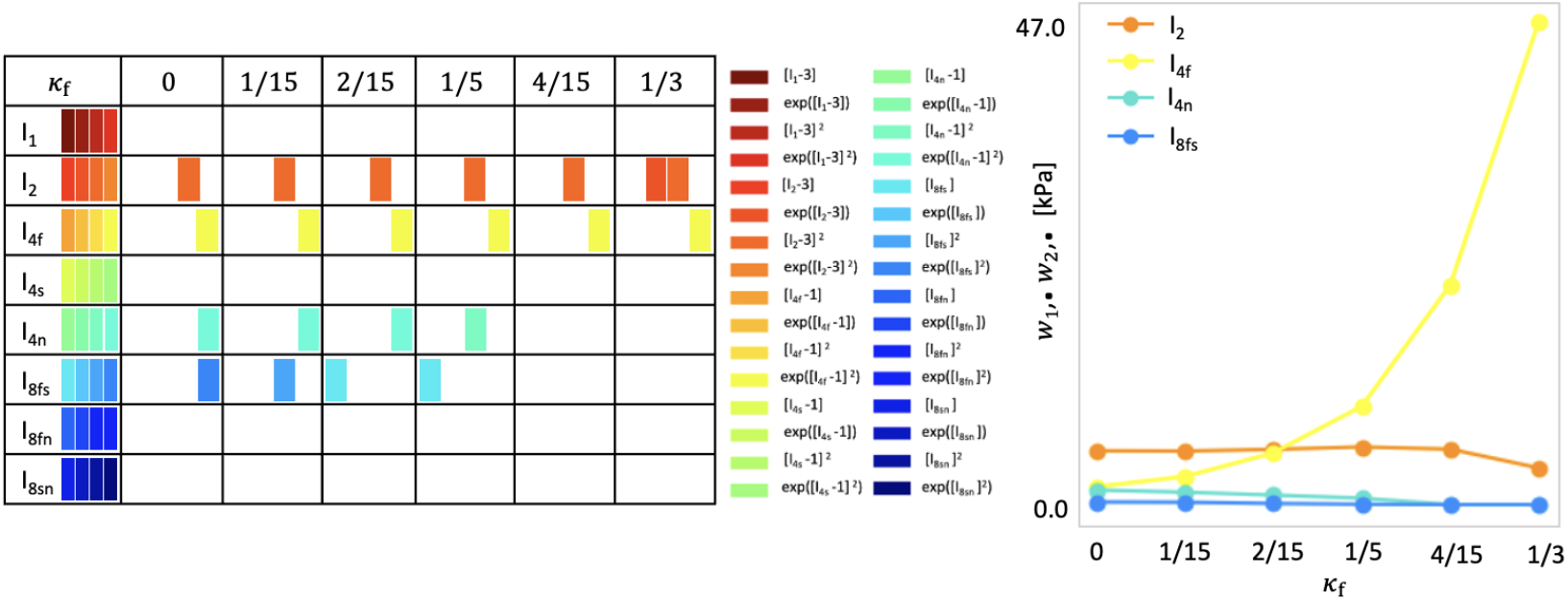
Model discovery for human myocardium with with varying amount of dispersion in fiber direction. Left: Active invariants for varying amount of dispersion in sheet and normal directions, *κ*_f_ ∈ {0, 1*/*15, 2*/*15, 1*/*5, 4*/*15, 1*/*3}, *κ*_s_ = *κ*_n_ = 0; color-coded blocks represent the contributions of the particular active term out of four possible terms, depending on one of the eight possible invariants given in the first column. Right: Stiffness-like weight values for active invariants; the sum of products 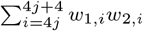, where *j* = 1, *j* = 2, *j* = 4, *j* = 5 for contributions of terms depending on *I*_2_, *I*_4f_, *I*_4n_, *I*_8fs_.

**Fig. 7.**
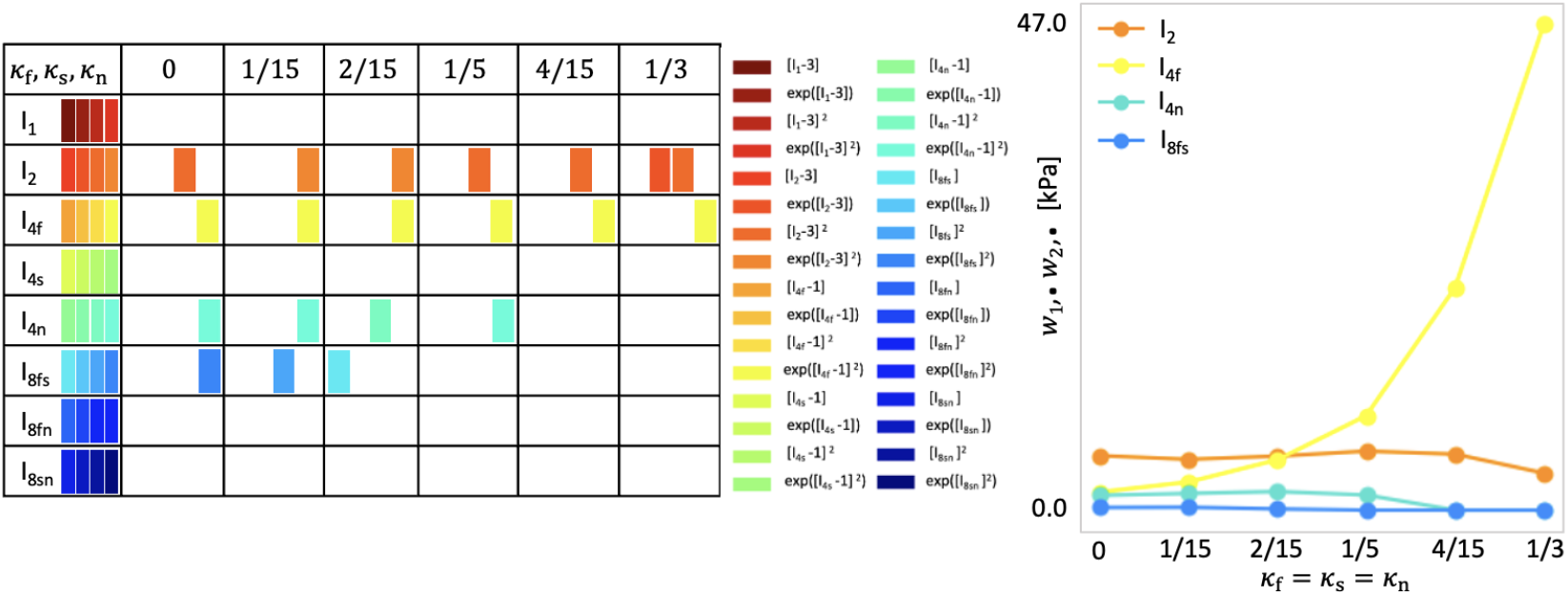
Model discovery for human myocardium with varying amount of dispersion in fiber, sheet and normal directions. Left: Active invariants for varying amount of dispersion in fiber, sheet and normal directions, *κ*_f_ = *κ*_s_ = *κ*_n_ ∈ {0, 1*/*15, 2*/*15, 1*/*5, 4*/*15, 1*/*3}; color-coded blocks represent the contributions of the particular active term out of four possible terms, depending on one of the eight possible invariants given in the first column. Right: Stiffness-like weight values for active invariants; the sum of products 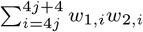, where *j* = 1, *j* = 2, *j* = 4, *j* = 5 for contributions of terms depending on *I*_2_, *I*_4f_, *I*_4n_, *I*_8fs_, respectively.

**Fig. 8.**
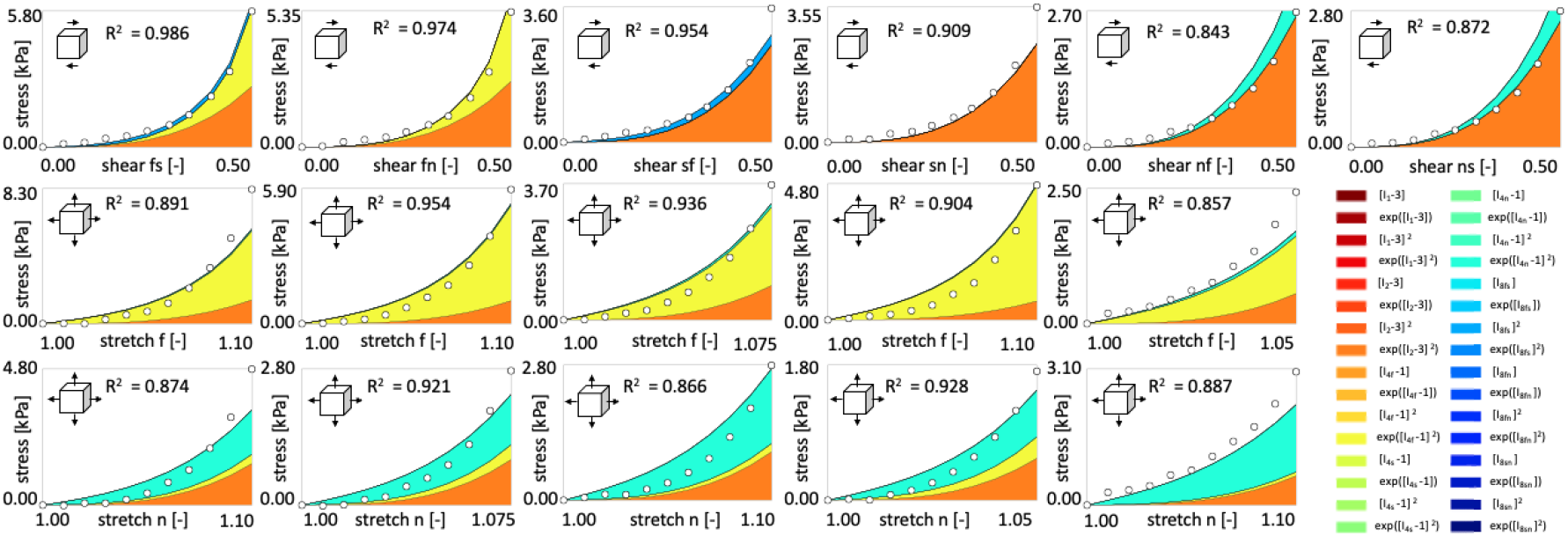
Human myocardial tissue data from triaxial shear and biaxial extension tests and the discovered model incorporating small dispersion in fiber, sheet and normal directions. First row displays Cauchy stress components as functions of shear strains during triaxial shear tests, second and third rows display stretches during biaxial extension tests in fiber and normal directions, respectively. Experimental data from (Sommer et al. 2015), illustrated with dots, are used for training the network from Figure 2 with dispersions in fiber, sheet and normal direction *κ*_f_ = *κ*_s_ = *κ*_n_ = 1*/*15. Color-coded areas highlight the activated terms, out of 32 possible terms, to the stress components, derived from the discovered free energy function *ψ*.

**Fig. 9.**
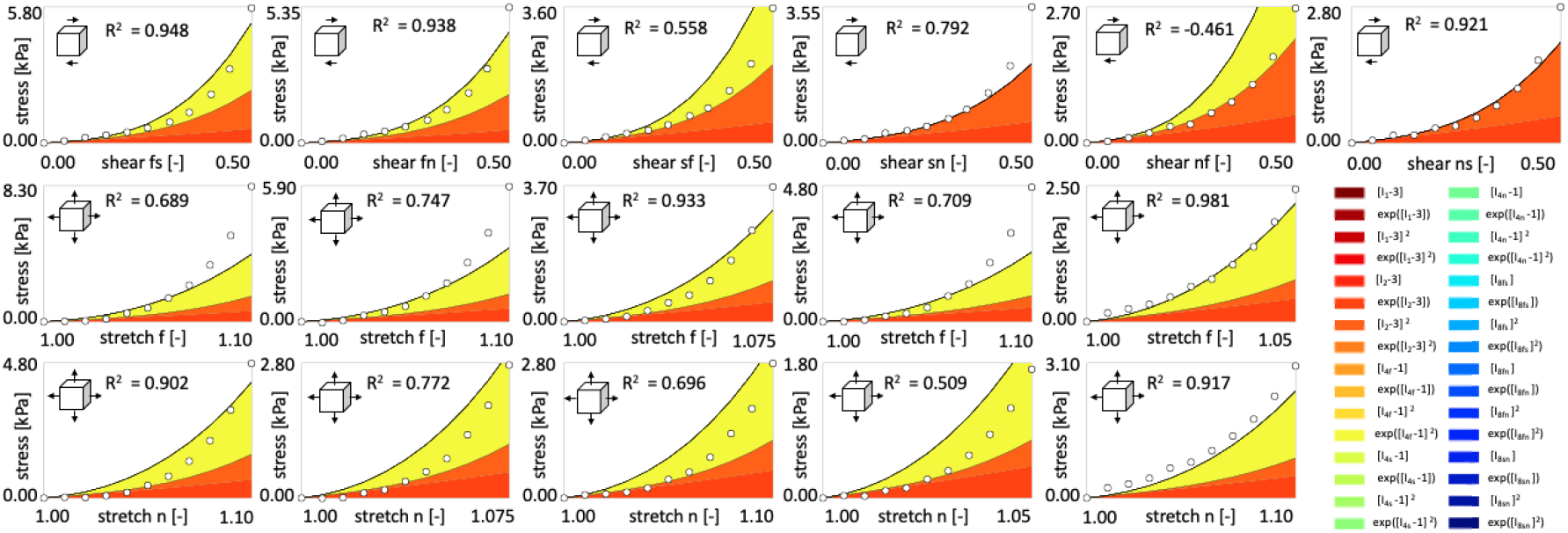
Human myocardial tissue data from triaxial shear and biaxial extension tests and the discovered model incorporating full dispersion in fiber, sheet and normal directions. First row displays Cauchy stress components as functions of shear strains during triaxial shear tests, second and third rows display stretches during biaxial extension tests in fiber and normal directions, respectively. Experimental data from (Sommer et al. 2015), illustrated with dots, are used for training the network from Figure 2 with dispersions in fiber, sheet and normal direction *κ*_f_ = *κ*_s_ = *κ*_n_ = 1*/*3. Color-coded areas highlight the activated terms, out of 32 possible terms, to the stress components, derived from the discovered free energy function *ψ*.

### Comparison to fitted dispersion in Holzapfel Ogden model

When calibrating the Holzapfel Ogden model with dispersed invariants 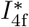 and 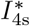, a study reports dispersion parameters *κ*_f_ = 0.00765 and *κ*_s_ = 0.0249 (Eriksson et al. 2013). Using this set of dispersion parameters for our model discovery, we recover the same four-term model depending on the invariants 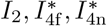 and *I*_8fs_. The goodness of fit of R^2^ = 0.896 increases slightly compared to the non-dispersed network with R^2^ = 0.890, but it is still lower than for some particular settings of *κ*_*i*_, *i* ∈ {f, s, n}, see Tables 1, 2 and 3. In particular, the most suitable dispersion parameters in term of the goodness of fit are *k*_*f*_ = 0 and *k*_*s*_ = *k*_*n*_ = 4*/*15 resulting in R^2^ = 0.922. This indicates that assuming a perfect fiber alignment and, more importantly, adding a dispersion in the normal direction, significantly improves the goodness of fit.

### Limitations and Outlook

First, our current model assumes full incompressibility, which may not fully represent the mechanical behavior of myocardial tissue that exhibits slight compressibility (Bonnemains et al. 2018; Avazmohammadi et al. 2020). Future research could explore a nearly incompressible formulation by incorporating an additional trainable parameter representing the bulk modulus. Second, fiber dispersion is represented exclusively through the fourth invariant, which may limit the model’s ability to capture more complex anisotropic effects. Although we lack of experimental data regarding the dispersion in the coupling invariants, future modification of the coupling invariants is possible (Melnik et al. 2018). Third, our training was conducted on a limited dataset. Efforts to obtain or simulate broader datasets would enhance training robustness. Finally, converting the dispersion parameter *κ*_*i*_ into a trainable parameter could improve the flexibility and precision of the model in capturing the responses of diverse materials.

## 4 Conclusion

In this work, we investigated the effects of fiber, sheet, and normal dispersions and random noise on the measured data in passive material model discovery using an orthotropic perfectly incompressible constitutive neural network for human myocardium. Overall, despite the non-convex nature of the minimization problem, our approach consistently identifies similar material models. This result underscores the robustness and reliability of the automated model discovery. Added random noise up to 7% has no influence on the discovered model. A small dispersion in the fiber direction of 0 *< κ*_*f*_ *≤* 2*/*15, an arbitrary dispersion in the sheet and normal directions of *κ*_s_ = *κ*_s_ *>* 0 with perfectly aligned fibers (*κ*_f_ = 0), or a small equal dispersion of *κ*_f_ = *κ*_s_ = *κ*_s_ = 1*/*15 all improve the goodness of fit and recover our previously discovered four-term model, subject to the isotropic invariant *I*_2_, the dispersed fourth invariants 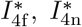, and the coupling invariant *I*_8fs_ in their quadratic or quadratic exponential forms. Introducing a constitutive neural network that accounts for dispersion in three directions enables us to account for the micro-structural architecture, to discover the best dispersion parameters, and to interpret model sensitivity with respect to the fiber, sheet, and normal directions.

## Supplementary information

Our source code, data, and examples are available at https://github.com/LivingMatterLab/CANN.

## Acknowledgments

This work is supported by the ERC Advanced Grant 101141626 Discover to Ellen Kuhl and Denisa Martonová, and by the NSF CMMI Award 2320933 Automated Model Discovery for Soft Matter to Ellen Kuhl.

